# The Shape of Biological Metadata: Measuring Repository Richness with Entity-Based NLP Metrics

**DOI:** 10.64898/2026.09.08.749906

**Authors:** Maria Juliana Rodriguez-Cubillos, Tomasz Zieliński, Jason R. Swedlow, T. Ian Simpson, Andrew J. Millar

## Abstract

Ensuring the availability and accessibility of research data is fundamental to advancing knowledge, as codified in the FAIR principles (Findable, Accessible, Interoperable, and Reusable). Accurate **metadata** documentation is indispensable for meeting these principles; however, entries in deposition databases often contain inadequate, repetitive, or incomplete descriptions.

Much of this metadata is captured in free-text fields, motivating the need for scalable, repository-agnostic methods to quantify metadata richness. Here, we quantify free-text metadata richness across three repositories using Natural Language Processing (NLP) methods: BioDare2, an experimental circadian rhythm database; DataShare, a domain-agnostic University of Edinburgh database; and Image Data Resource (IDR), a public repository of biological image datasets from published studies.

In general, repositories exhibit distinct distributions of word count and information density, consistent with differences in scope. Named-entity recognition and information-density metrics detected significant category-level differences in BioDare2 (species) and DataShare (communities), while identifying greater consistency in the more curated IDR. The most frequent entities reflected each repository’s focus: circadian terminology in BioDare2, microscopy-related entities in IDR, and community-driven terms in DataShare.

We develop a scalable framework utilising word counts, named-entity recognition, and entity-derived information density to assess metadata quality across repositories, offering a broadly applicable evaluation tool.

## 1 Introduction

### 1.1 Metadata: A Foundational Overview

Metadata is created to describe the underlying data in a detailed and unambiguous way by providing context, recording transactions, reporting content, and situating the data Greenberg, Wu, and Liu (2023); Ulrich et al. (2022). It is essential for finding, retrieving, and reusing research outputs. Providing relevant scientific data requires high-quality metadata to ensure that appropriate datasets can be discovered and reused effectively Gonçalves, O’Connor, Martínez-Romero, Graybeal, and Musen (2017); Wu et al. (2023).

Interest in metadata quality has grown over the last few decades due to the rapid development of computational tools for managing digital information, data-sharing policies, interoperability goals, and open-access efforts across different scientific communities worldwide Greenberg et al. (2023). Moreover, this landscape has changed swiftly due to the global adoption of data management policies such as the FAIR principles, which seek to make digital datasets Findable, Accessible, Interoperable and Reusable Wilkinson et al. (2016).

Despite the efforts to promote and adopt these principles for the past ten years, multiple empirical studies have shown a clear need to improve data and metadata quality Ugochukwu and Phillips (2024); Zaveri, Hu, and Dumontier (2019). Software systems that index and utilise data can fail to retrieve otherwise suitable results for lack of metadata Gonçalves and Musen (2019). To solve that, many public repositories offer user-friendly tools and interfaces for entering and querying metadata. Nevertheless, researchers are often not constrained to use standard terminology or fail to provide accurate and detailed information Assante, Candela, Castelli, and Tani (2016), as the majority of the metadata comes as free text.

In biological sciences, the spread of automated experimental platforms is emphasising a dichotomy between biological and technical metadata. The researcher must usually identify themselves and manually specify the identity of the biological samples (perhaps via an intermediate sample management system). The platform can automatically write the internal settings that were used in a study into its digital outputs, such as the objective lens and illumination for an image from an automated microscope.

### 1.2 NLP approach to analyse metadata

Recent breakthroughs in artificial intelligence (AI) have enabled a rich environment for improving and suggesting metadata to make it more FAIR, such as Natural Language Processing (NLP) methods. Therefore, NLP focuses on natural language translation, information retrieval and extraction, text summarisation, question answering, topic modelling, and opinion mining Chowdhary (2020). Text mining is one of the first activities linked solidly to those tasks, which capture implicit knowledge from textual data Jo (2019). Nevertheless, datasets are not ‘AI-ready’ if the metadata is insufficient Sundaram and Musen (2023).

Automated NLP can be used to examine data descriptions effectively and accurately as a human does, specifically for a limited amount of text Chowdhary (2020). Different packages have been coded to efficiently perform some NLP and text mining tasks. Open-source tools like NLTK Bird, Klein, and Loper (2009) and SpaCy Honnibal and Montani (2017) have recently fostered NLP applications for scientific communities. For example, SpaCy can recognise the nature of words in a sentence and their dependence on the other words using a combination of rules-based and statistical methods Bird et al. (2009); Blanchy, Albrecht, Koestel, and Garré (2023); Honnibal and Montani (2017); Loper and Bird (2002).

In the repositories examined here, researchers are required to complete submission forms with multiple fields. Some fields are structured (e.g., predefined response formats or multiple-choice items), with allowable options set by database managers in advance. However, purpose, aims and experimental details primarily necessary for data aggregation or meta-analysis are typically provided in free-text, natural language fields. Therefore, our study analyses metadata contained in unstructured text, a format commonly used in public repositories.

### 1.3 Biological repositories studied

The analysis focused on metadata from three repositories: BioDare2 Zielinski, Hay, and Millar (2022), DataShare of Edinburgh (2008); Rice and Haywood (2011), and the Image Data Resource (IDR) Williams et al. (2017). The first repository, BioDare2, is a free online service that hosts research data on 24-hour biological rhythms, termed circadian rhythms. It also provides rapid time-series analysis, summary statistics and visualisations, which researchers use routinely to analyse unpublished data from ongoing research. Access is granted on the condition that deposited data will be made public after three years Zielinski et al. (2022).

BioDare2 stores metadata in a relatively informal and less detailed structure, comprising free-text descriptions, species information, authorship, and techniques Zielinski et al. (2022). This service has gained international popularity, accumulating over 20,471 records as of July 2025, and has been used to test rhythms in more than 39 species. Within BioDare2, users can select the species name relevant to their studies from a dropdown menu, enabling the classification and comparison of data across different groups. The global usage of BioDare2 is illustrated in Supplementary Figure 1 - S1, which displays the number of users from January to November 2023. Due to the absence of a dedicated API, metadata were obtained as JSON files exported by the database manager. Notably, the platform lacks a formal curation process, with only minimal length requirements imposed on certain fields.

The secondary repository, DataShare, a subject-agnostic database and the University of Edinburgh’s digital deposition repository was included. Research data are divided into five communities associated with different Colleges of the university, as well as thematic collections and support services. This institutional repository was established to allow researchers to publicly disseminate datasets as one of the research outcomes. In contrast with BioDare2, DataShare is considered semi-curated, as deposits are manually evaluated for relevance to the repository scope, valid layout and format, and exclusion of spam. The repository curators also offer suggestions for enhancing metadata quality by email of Edinburgh (2008).

Finally, The Image Data Resource (IDR) is a public repository of biological image datasets from published scientific studies, enabling the community to submit, search, and access high-quality biological image data. It links information from various imaging methods, such as multi-dimensional microscopy and digital pathology, with public genetic or chemical databases, and cell and tissue phenotypes expressed using controlled ontologies. Because of that, it provides the landscape to analyse gene networks and show functional integration that is inaccessible to individual studies Williams et al. (2017). The records in this database are linked to published studies, ensuring that the information corresponds to completed and reviewed experiments, rather than preliminary data.

This study aims to quantify metadata richness across these three public repositories to establish a baseline of available information, evaluate the effectiveness of an entity-recognition–based analysis, and identify metrics to improve metadata in these repositories.

## 2 Methodology

### 2.1 Metadata retrieval

From BioDare2, 20,471 entries were provided for analysis, taken from the creation of the database up to June 2025. The database submission form includes four free-text fields: “Name” (minimum 55 characters) and “Purpose” (minimum five words), which are mandatory, and “Comments” and “Description”, which are optional. In addition, users must specify “Species”, selected from 39 options in a drop-down menu, and “Data category”, selected from 12 options in a second drop-down menu. The “Contributors” field is automatically populated with user information. All these fields were included in the data-retrieval code, along with the submission date.

For DataShare, metadata were retrieved via the REST API (https://datashare.ed.ac.uk/rest). In total, 7,754 records were harvested, covering the period from the repository’s launch to June 2025. Among the available metadata fields, only the “Abstract description” field was analysed, as it was the only free-text field consistently present across records and exhibited sufficient variation to support comparative analysis.

For IDR, metadata were obtained from GitHub, where the repository stores all associated metadata (https://github.com/IDR/idr-metadata). In total, 132 entries were retrieved, covering the full database up to June 2025. As with DataShare, only the “Study description” and “Experiment description” fields were analysed, as they were the only free-text fields consistently available across all entries included in the analysis. IDR is a curated resource that organises data using structured fields, displayed as key-value pairs. Since descriptions are free-text fields, they provide a natural and fair basis for comparison with other resources.

### 2.2 Analysis methods

The code used to analyse the repositories was written in Python (available here https://github.com/mjrodriguezc/metadata_project) and designed to be easily customised to all three platforms. It converts the JSON files into a single data frame per repository using the Pandas package, while preserving a consistent column structure. For all repositories, only free-text description fields were analysed. Nevertheless, additional variables (e.g., species and community) were used to support more granular comparisons within subsets of the data. As shown in Figure 1 summarises the overall workflow used to analyse and characterise the metadata.

**Figure 1:**
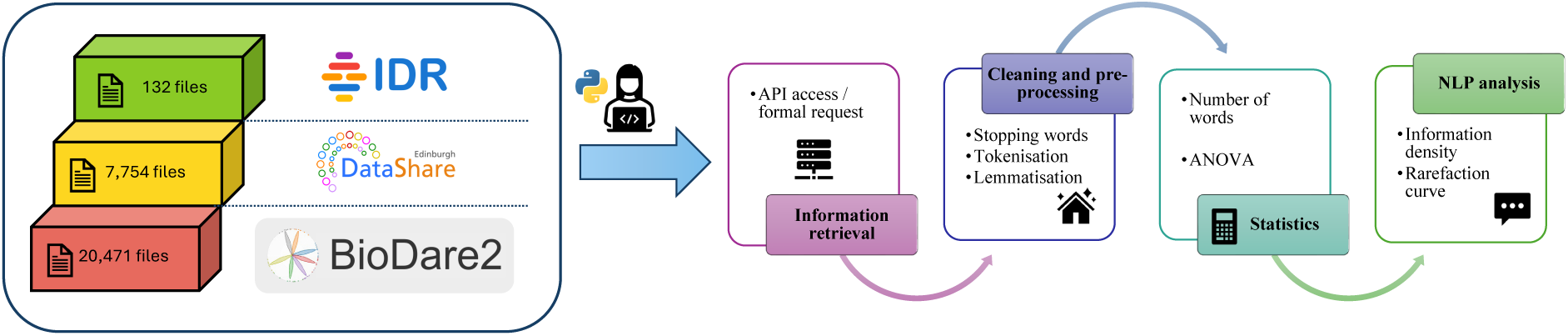
Workflow for the descriptive analysis of the repositories. It integrates the pre-processing steps with the NLP approach for entity recognition and explains the two sets of comparisons that were run to analyse the entire dataset. Three independent datasets were used during the study: BioDare2 with 20,471 entries, DataShare with 7,754 entries and IDR with 132 entries. Each one was processed separately, but the same processing steps were applied to each. First, the files were acquired from the repositories. Then, the stop words and special characters were removed, all capital letters were replaced with lowercase, and the words were lemmatised. Afterwards, the number of words was calculated, the entities were recognised in the text, and a one-way ANOVA (Analysis of Variance) were performed to identify differences between groups inside the repositories.

### 2.3 NLP analysis

The text was cleaned by removing stop words (e.g., conjunctions) and special characters, tokenise and lemmatised using the NLTK package Jo (2019); Loper and Bird (2002) to reduce lexical redundancy. Lemmatisation maps words that share a common root to a single canonical form (lemma), thereby facilitating entity recognition. For example, the verb “test” may appear as “testing” or “tested”; lemmatisation reduces these variants to the lemma “test”.

Subsequently, we quantify the number of words and characters in each description field. These measures were used to examine whether longer descriptions are associated with larger datasets or specific species, for example. For each experiment, word counts were summed across the selected fields.

Thereafter, we used SciSpaCy’s “en_core_sci_sm” model to identify named entities in the free-text descriptions Neumann, King, Beltagy, and Ammar (2019). This model is tailored to biomedical text Honnibal and Montani (2017) and is trained on resources such as the GENIA 1.0 Treebank Mcclosky and Charniak (2008) and word2vec embeddings that were, in turn, trained on the PubMed Central Open Access Subset Pyysalo, Ginter, Moen, Salakoski, and Ananiadou (2013).

To estimate information density, we divided the number of entities extracted from each entry by the corresponding total world count. This metric was computed on an entry-by-entry basis for each repository. Word clouds were generated in Python using the “wordcloud” package Mueller (2015). Finally, for entity quantification, we divided the information into existing set categories within the datasets. For the biological repositories, we utilized species names, whereas for DataShare, we used community names. We then removed the 10 most frequent entities from each repository to better capture terms that were more specific to each subset.

## 3 Results

### 3.1 Patterns in word count distribution differ between repositories and subsets

We quantified the text length per entry in each repository by counting the total number of words in the free-text description fields (Figure 2a). Overall, although BioDare2 is the largest of the three repositories by number of entries, it has the lowest word count and shows substantial variability (mean words count M = 18.0, SD = 20.07). When the BioDare2 dataset is stratified by species (Figure 2b), the word count per category does not differ substantially across groups, except for the Synthetic data category. This difference likely reflects metadata heterogeneity within this category.

**Figure 2:**
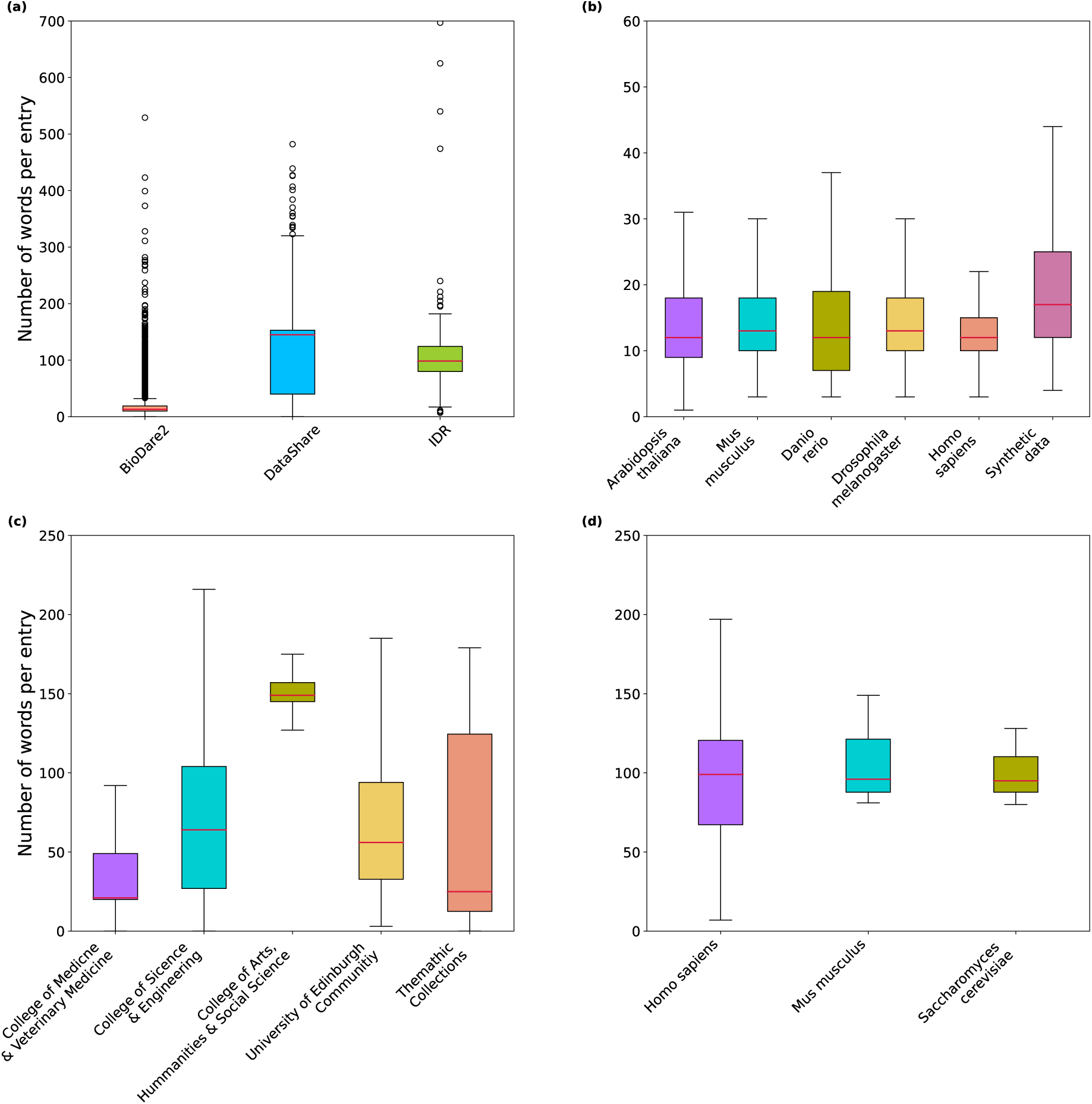
Word count per entry across repositories and subsets. (a) Distribution of the number of words per entry in BioDare2 (orange), DataShare (light blue), and IDR (green). The points falling outside the expected range are outliers within the datasets. Box is IQR whiskers normally +/- 1.5IQR. (b) Distribution of the number of words per entry in BioDare2 for the six most-studied species. (c) Distribution of the number of words per entry in DataShare for the five top communities. (d) Distribution of the number of words per entry in IDR for the three most-studied species (with more than 10 entries each). Outliers are not shown in panels (b–d).

For DataShare (Figure 2a), the large difference between the smallest and largest word counts may reflect the diversity of datasets hosted in the repository (M = 110.14, SD = 64.49). When the content is stratified into five categories (Figure 2c), the *College of Arts, Humanities and Social Sciences* community shows a substantially higher word count than the others. This community includes several large entries from the same collection with similar descriptions, which may account for the upward shift in the mean in the comparative box plot (Figure 2a) and help explain the differences observed in Figure 2c.

Moreover, it is crucial to note that subsets of entries with repeated descriptions can alter metrics such as mean word count within a repository, as observed for DataShare and the *College of Arts, Humanities & Social Science*. Although such repetition may indicate consistency across entries, it can also reduce descriptiveness and limit entity enrichment at the individual entry level.

For IDR (Figure 2a), the datasets are relatively consistent, and all are associated with published articles, which ensures a baseline level of curation for the deposited information (M = 113, SD = 95). The database focuses on microscopy images, and the observed variation is likely driven mainly by differences among species-specific datasets rather than by substantial diversity in data types, as seen in DataShare. In contrast, when the information in IDR was stratified by the three most frequently studied species, studies involving *H. sapiens* had longer descriptions in terms of word count than those for the other two species.

### 3.2 Information density across repositories showed similar behaviour between repositories with different levels of curation

We applied an NLP approach to identify and count named entities in the descriptions. For that purpose, we defined the metadata’s information density as the ratio of entities per word (Figure 3). A higher proportion of entities indicates a richer data description among entries with equal word count.

**Figure 3:**
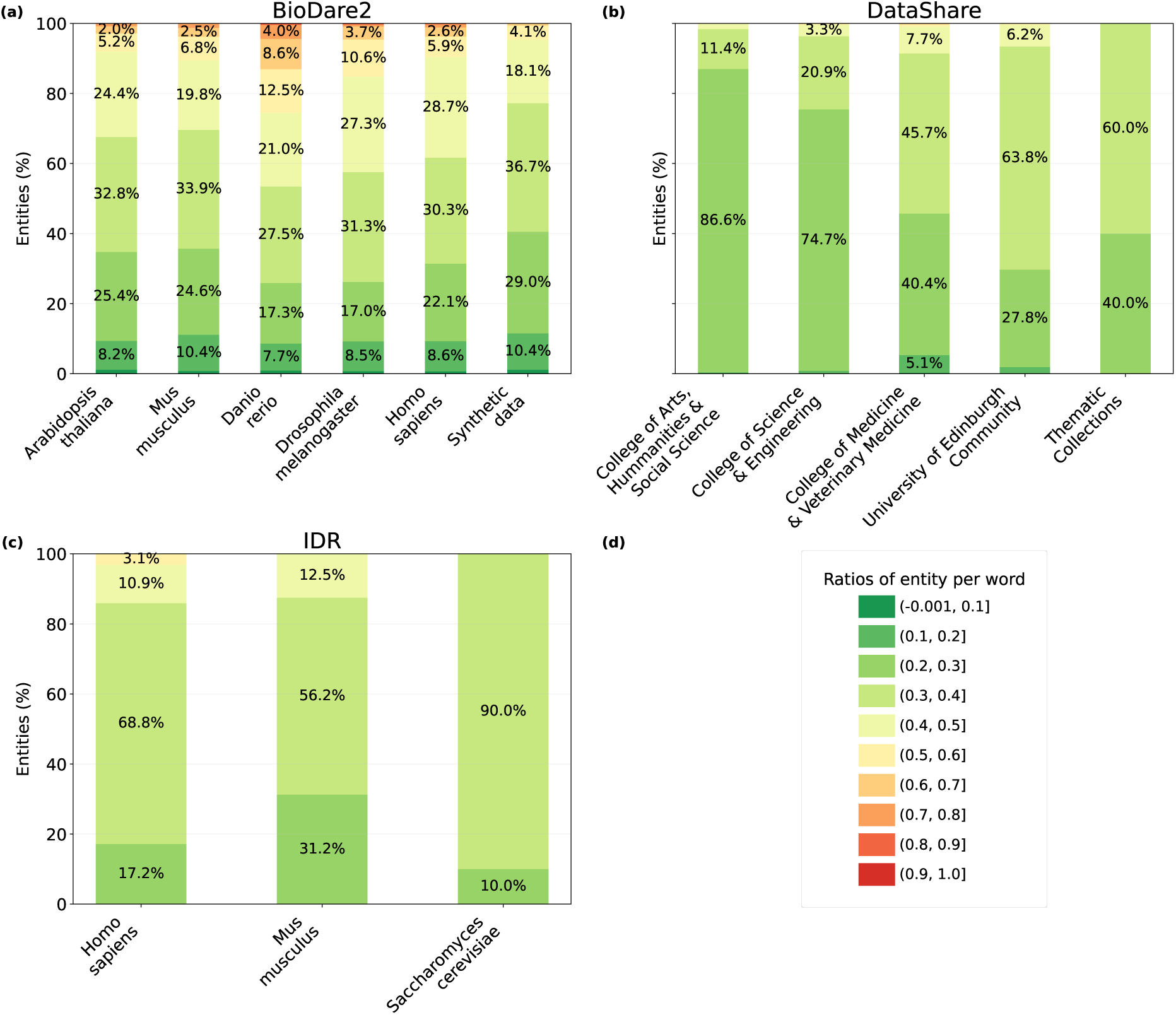
Information density across repositories and subsets. The distribution of information density is shown as the percentage of entries with a given entities-per-word ratio in (a) BioDare2 for the six most-studied species, (b) DataShare, for the five largest communities, and (c) IDR, for the three most-studied values, which were grouped into ten bins to facilitate visualisation.

Some descriptions were very information-dense, with up to 0.8 entities per word, though the plots show a broad distribution across the ten possible density bins for each data category. Nevertheless, a longer description is not necessarily more informative if the number of entities is low. For example, among *A. thaliana* entries in BioDare2, most descriptions (32.8%) contain between 0.3 and 0.4 entities per word (Figure 3a). The mean information density was 0.36 for descriptions from each species in BioDare2 with an SD of 0.13.

To test for differences in mean information density across categories, we conducted a one-way ANOVA. For BioDare2, the analysis showed a significant effect of category, F=92.2 with p=4.06557*10-95. In other words, there is strong statistical evidence that the mean information density differs across categories. For DataShare, a one-way ANOVA across data categories yielded an F-value of 119.921 with a p-value of 3.004*10-99, providing strong evidence that mean information density differs among categories. For IDR, a one-way ANOVA across the three most represented species yielded an F-value of 1.45 with a p-value of 0.2391. The between-group differences were therefore not sufficient to reject the null hypothesis of equal means; consequently, no post hoc test was performed. This indicates that the entities per word ratio does not differ significantly among the groups.

In DataShare and IDR, most values lie at or below 0.4 entities per word, with mean information densities of M = 0.029, SD = 0.045 and M = 0.34, SD = 0.05, respectively. The information densities significantly differed between descriptions from all 6 species in BioDare2, except *A. thaliana* and *M. musculus*, and between descriptions from the College of Arts, Humanities and Social Science and most other categories in DataShare, but not between the species in IDR (Table 1).

**Table 1:**
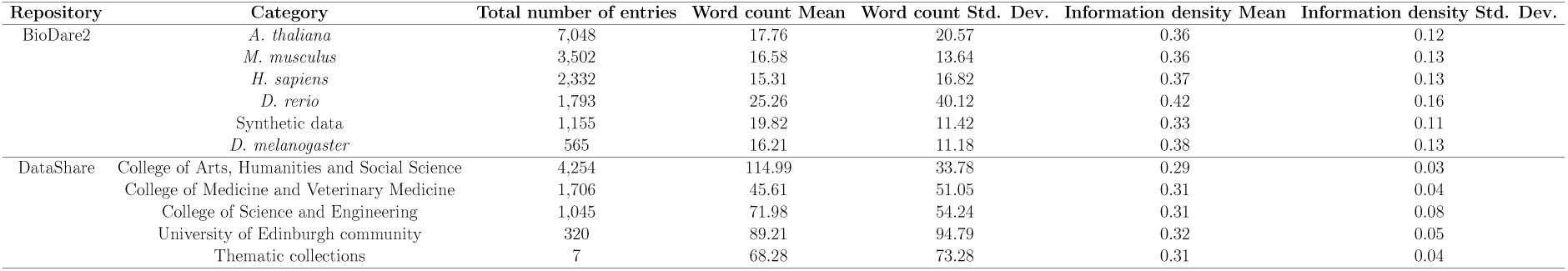
Comparison of word counts and information density between BioDare2 and DataShare datasets. The results show varying word count means and standard deviations across different categories within each dataset, with BioDare2 ranging from 15.31 to 25.26 and DataShare from 45.61 to 114.99. The information density means also differs, with BioDare2 ranging from 0.33 to 0.42 and DataShare from 0.29 to 0.32. Overall, the table highlights notable differences in linguistic characteristics between the two datasets, which may have implications for text data analysis and interpretation.

| Repository | Category | Total number of entries | Word count Mean | Word count Std. Dev. | Information density Mean | Information density Std. Dev. |
| --- | --- | --- | --- | --- | --- | --- |
| BioDare2 | *A. thaliana* | 7,048 | 17.76 | 20.57 | 0.36 | 0.12 |
|  | *M. musculus* | 3,502 | 16.58 | 13.64 | 0.36 | 0.13 |
|  | *H. sapiens* | 2,332 | 15.31 | 16.82 | 0.37 | 0.13 |
|  | *D. rerio* | 1,793 | 25.26 | 40.12 | 0.42 | 0.16 |
|  | Synthetic data | 1,155 | 19.82 | 11.42 | 0.33 | 0.11 |
|  | *D. melanogaster* | 565 | 16.21 | 11.18 | 0.38 | 0.13 |
| DataShare | College of Arts, Humanities and Social Science | 4,254 | 114.99 | 33.78 | 0.29 | 0.03 |
|  | College of Medicine and Veterinary Medicine | 1,706 | 45.61 | 51.05 | 0.31 | 0.04 |
|  | College of Science and Engineering | 1,045 | 71.98 | 54.24 | 0.31 | 0.08 |
|  | University of Edinburgh community | 320 | 89.21 | 94.79 | 0.32 | 0.05 |
|  | Thematic collections | 7 | 68.28 | 73.28 | 0.31 | 0.04 |

### 3.3 Rarefaction Curves as a Tool for Informing Metadata Development in Repositories

To determine whether each entire repository is saturated with respect to named-entity diversity—or whether it could still expand as additional entries are deposited—we generated rarefaction curves showing the cumulative number of unique entities as a function of the number of entries across datasets Hurlbert (1971). Given the substantial differences in the number of entries in BioDare2, DataShare, and IDR, the comparisons are most appropriate between the two largest repositories (Figure 4a). We also performed the analysis per metadata categories in BioDare2 (Figure 4b) and DataShare (Figure 4c). Notably, neither curve reaches a plateau, indicating that additional entries would likely continue to introduce new entities in both repositories.

**Figure 4:**
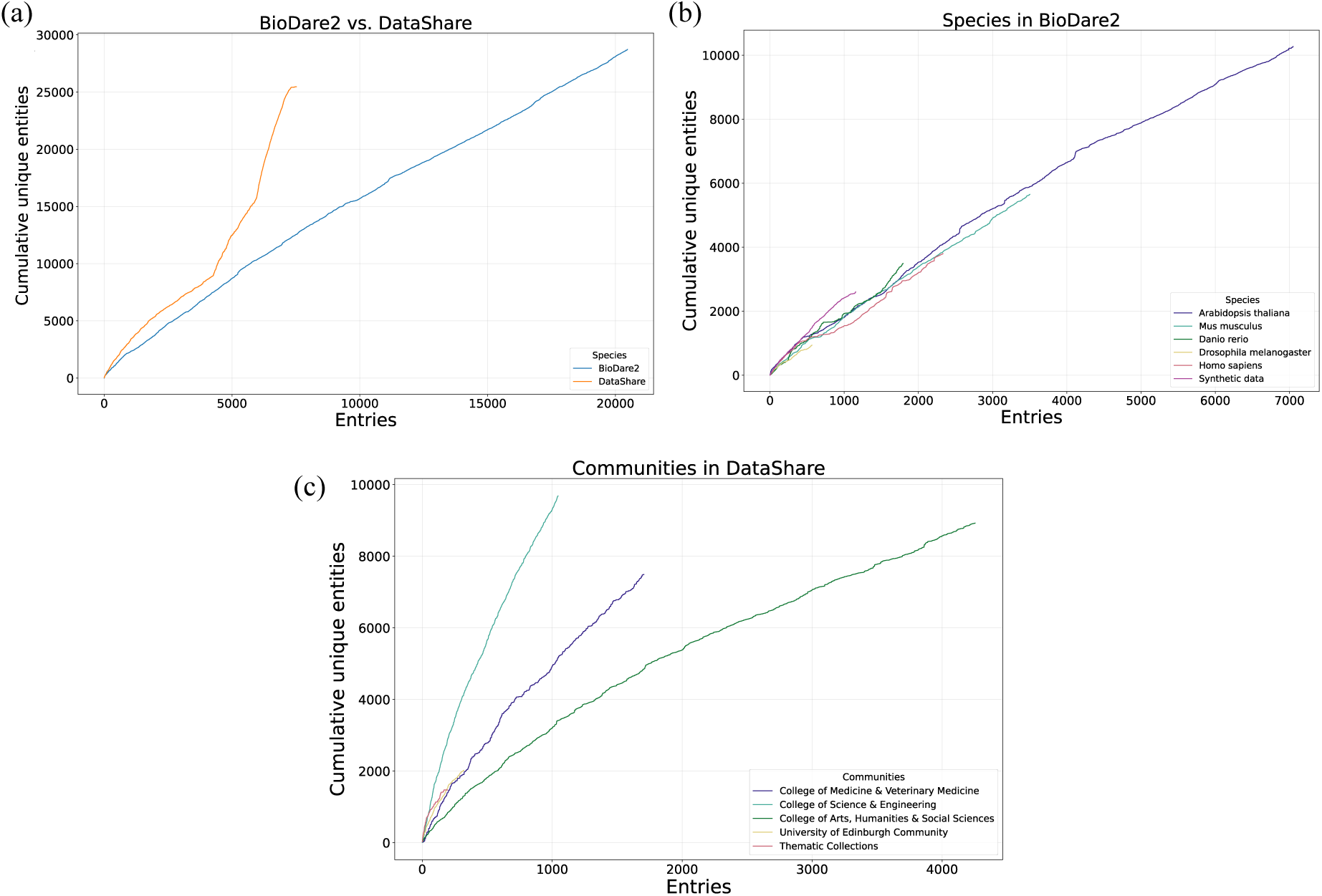
Rarefaction analysis of entity discovery with increasing entries. Rarefaction curves show the relationship between the number of entries sampled and the cumulative number of unique entities identified in (a) BioDare2 (light blue) and DataShare (orange). The same analysis is shown for (b) BioDare2 across its six most-studied species or data types, and for (c) DataShare across its five largest communities. None of the curves reaches a plateau, indicating that additional entries are likely to reveal further entities and that these databases have not yet captured the full diversity of entity information.

When rarefaction curves were calculated by species for BioDare2 (Figure 4b) and by the largest communities for DataShare (Figure 4c), none of the categories reached saturation, indicating that this pattern is consistent across groups. We also performed the same analysis for IDR, where the trend persisted (Supplementary Figure 2). Please note that this analysis is not time-oriented, and the ordering of entries does not follow a temporal sequence.

We used word clouds to highlight entities that recur across entries, with larger terms indicating higher frequency in each repository (Figure 5). Even when this technique could be useful to extract meaningful information, given the nature of the analysis and the model used, some domain-specific abbreviations or incorrectly handled gene symbols may not be apparent in this representation.

**Figure 5:**
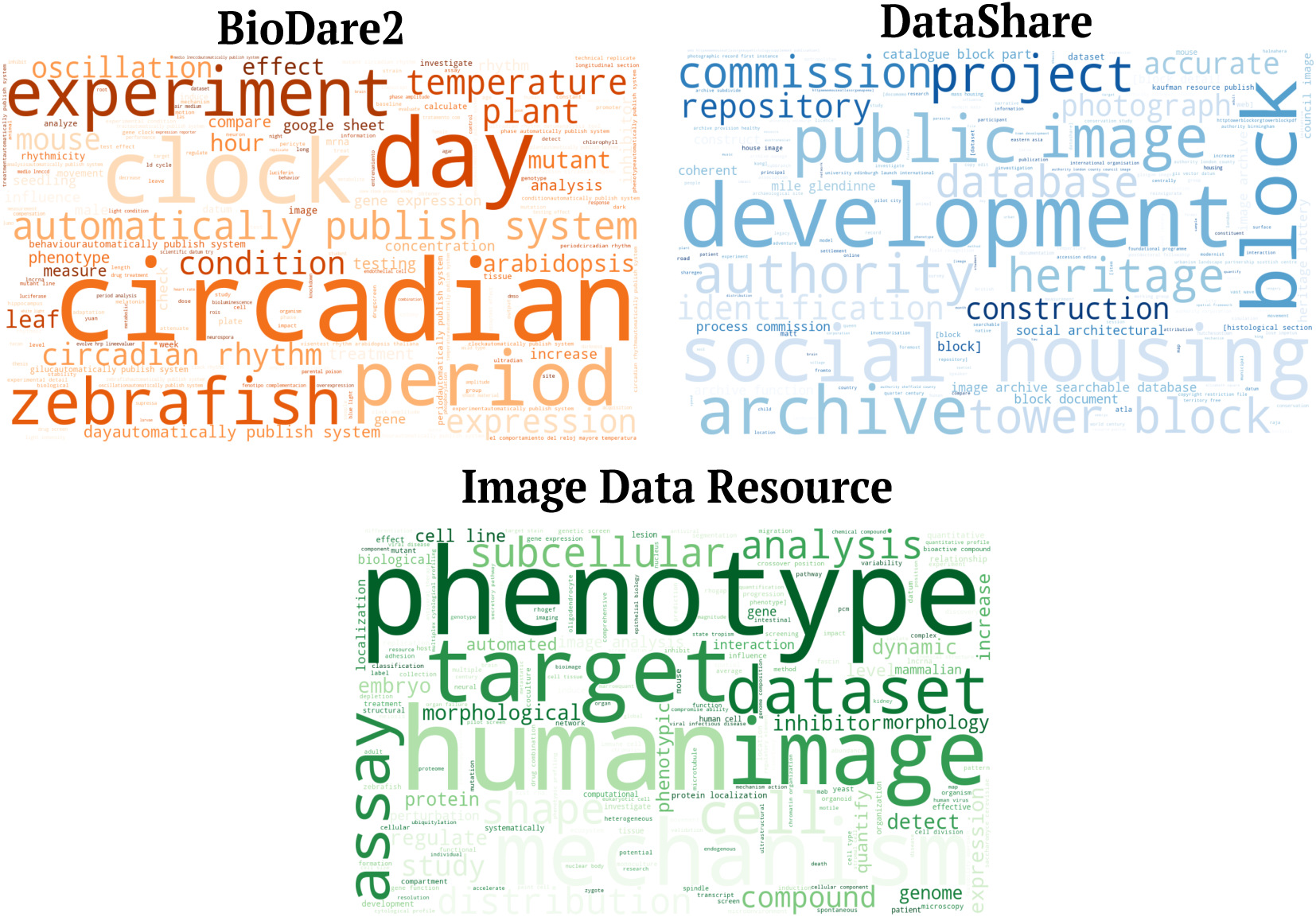
Word clouds of term frequency across repositories. Word clouds for BioDare2 (orange), DataShare (blue), and IDR (green). Word size is proportional to term frequency in each dataset: larger words indicate terms that occur more often.

In BioDare2, “circadian”, “day”, and “experiment” reflect information related to circadian rhythms. In DataShare, “block”, “development”, and “project” are driven by the largest community in the repository, which is related to high-rise housing. Finally, in IDR, “phenotype”, “human”, and “image” capture the repository’s focus on microscopy images.

Finally, we examined this pattern across four subsets in BioDare2 to determine whether the extracted entities also capture species-specific experimental context or biological features (Figure 6 6). To increase the granularity of the analysis, we removed the 10 most frequent entities in the full dataset and then plotted the 15 most frequent named entities within each subset. In this analysis, all entities were manually classified into four broad categories: genetic elements, experimental conditions, the phenotype studied, and organism condition. Among the most salient entities identified, the *A. thaliana* dataset included “Arabidopsis”, “seedling”, “leaf”, “tissue”, and “movement” as representative terms (Figure 6a),. The last of these likely relates to leaf movement rhythms. Measuring these movements provides a simple, reliable assay for the plant circadian clock and, unlike systems based on the firefly luciferase reporter gene, requires no prior genetic manipulation of the plant Edwards and Millar (2007).

**Figure 6:**
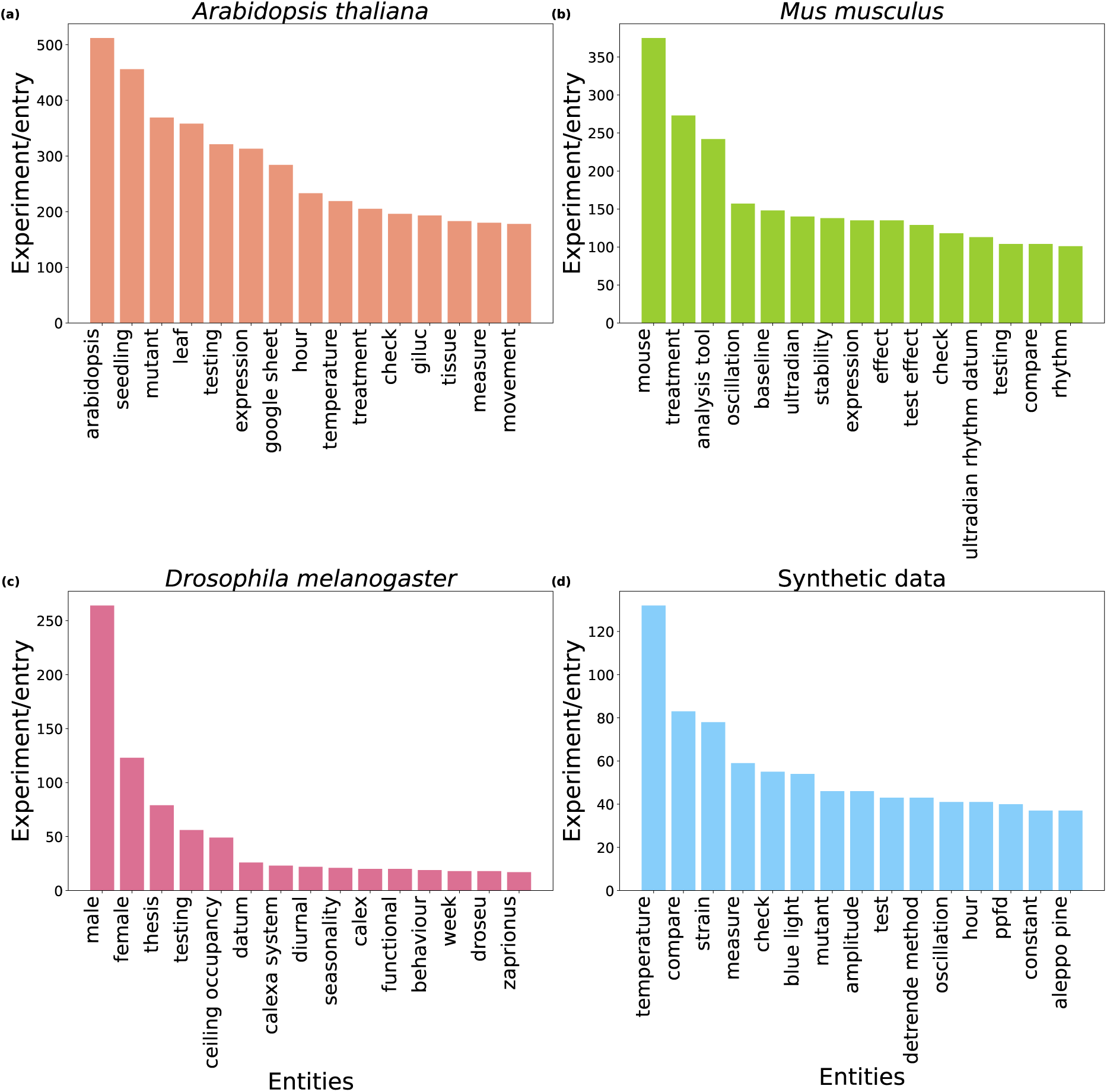
Most frequent subset-enriched entities in BioDare2. Number of occurrences of the 15 most frequent named entities within each subset after removing the 10 most common entities in BioDare2 for (a) *A. thaliana*, (b) *M. musculus*, (c) *D. melanogaster*, and (d) Synthetic data. To highlight entities that are characteristic of each subset (rather than globally frequent across BioDare2), the 10 most common entities in the full database were excluded before selecting the top 10 entities for each subset.

Representative entities in the *M. musculus* dataset (Figure 6b) include “mouse”, as expected, as well as “ultradian” and “oscillation”. The term “ultradian” denotes rhythms with periods ranging from fractions of an hour to several hours, which affect many processes including the same biological markers as circadian rhythms Copinschi and Challet (2016). BioDare2 users can adapt the platform’s tools to analyse ultradian rhythms Zielinski et al. (2022).

For *D. melanogaster* (Figure 6c), “male”, “female”, “behaviour” and “calexa system” entities are easily related to the organism. For instance, CaLexA (calcium-dependent nuclear import of LexA) is a genetic technique employed in Drosophila melanogaster to label and visualise active cells in living insects, most notably neurons and enteroendocrine cells Masuyama, Zhang, Rao, and Wang (2012).

Overall, the extracted entities were coherent with the corresponding species. Surprisingly, when we analysed the “synthetic data” subset (Figure 6d), we did not identify terms indicative of modelling or a mathematical context, as would be expected for this category. Entities such as “ppfd” (photosynthetic photon flux density, a measure of light intensity), “temperature”, “strain”, and “aleppo pine” were unexpected (Figure 6d). Inspection of this subset found descriptions that mention a range of diverse plant species, confirming our suspicion that users may select the “synthetic data” option as a default when the organism they studied is not yet available in the species list provided by the submission form.

## 4 Discussion

The results showed how repositories differed in both text volume and information density, with significant category-level differences in BioDare2 and DataShare but not in IDR. Rarefaction curves did not plateau, indicating continued growth in unique entities with additional deposits. The distribution of frequent entities was consistent with repository scope and, in BioDare2, with species-specific experimental context, with the synthetic-data subset showing a distinct pattern.

Taken together, the word count, the word cloud, the information-density analysis, named-entity extraction and rarefaction curves provide a consistent view of how descriptive metadata varies across repositories and categories. We propose a series of calculations to evaluate metadata richness across different repositories. We start with simple measurements such as word count, which can reveal differences in behaviour among metadata subsets, and progress to more complex analyses, including rarefaction curves that illustrate the status of the entire repository.

Simple metrics such as word count can be surprisingly informative. Our analysis shows that a repository’s scope can influence the length of free-text metadata: general repositories, such as DataShare, tend to display greater heterogeneity Assante et al. (2016) and longer descriptions, whereas specialist repositories such as IDR provide a baseline level of curation and thus more consistent descriptions Williams et al. (2017), as we observed (Figure 2). However, metrics like word count may reflect not only depositor behaviour but also the repository’s overarching mission and community practices. Ultimately, the repository’s design and community practices together influence how depositors describe their data.

In this study, word clouds were primarily used to assess the alignment between each repository’s disciplinary context and the content of its individual entries, as determined by the entities identified through named entity recognition (NER). By displaying domain-relevant terms, word clouds provide qualitative evidence that NER is successfully extracting meaningful information from the database or about missing information, for instance, we found that some entries with comments sometimes pointing to other sources, like “details are in the Google sheet”, for more information.

The advantage of using NER lies in its ability to localise and classify text into predefined entity categories such as organisms, institutions, or locations Li, Sun, Han, and Li (2022). However, NER-derived entities offer only an approximate representation of the information present, as they are limited to what the NLP model can recognise. Furthermore, employing pretrained NLP models specifically trained on biological data, such as SciSpacy Neumann et al. (2019), enables the identification of a detailed number of entities within each text, as demonstrated in Figure 5. We also applied this approach to determine the most frequently mentioned entities within each data subset, thereby facilitating the linkage of information in the free-text descriptions to particular species, as shown in Figure 6.

For example, our analysis shows that BioDare2 entries, even when they contain shorter free-text descriptions, can produce multiple relevant entities (see Supplementary Figure 3 - S3). In fact, longer text may dilute the density of identifiable entities, whereas concise descriptions can remain entity-rich, as shown in Figure 3. Therefore, it is preferable to assess richness using information density alongside other metrics, rather than relying on a single measure for entry evaluation. This approach also benefits users, as it removes the need to provide a predefined list of keywords or demonstrate detailed semantic understanding.

However, we also acknowledge limitations inherent to NER-based methodologies. Specifically, measured richness is constrained by the recogniser’s coverage: highly domain-specific entities may be overlooked if the NLP model’s training corpus does not sufficiently represent relevant terminology Ritter, Clark, and Etzioni (2011). While our analysis successfully identified entities consistent with the biological context of the entries—particularly in the case of synthetic data in BioDare2—it is notable that no irrelevant terms were observed among the 11th to 25th most frequently identified entities. For those cases, the semantic context afforded by NER approaches may facilitate the identification of entities originating from outside the domain of biology, a prospect that warrants further investigation.

Entities can also be used to quantify the information richness of free-text descriptions. This is especially helpful when evaluating longer descriptions, as a higher word count does not necessarily indicate that more meaningful entities are present. Therefore, information density—the number of relevant entities per word in each text—is a useful metric for assessing the quality of information, as shown in Figure 3. It can be applied not only to individual entries but also as a parameter for comparing different data subsets.

Entity counts can also be analysed at a broader scale. Using rarefaction curves, we address two related questions: how much information the database contains in terms of unique named entities, and whether the existing content has captured most of the entities that the repository is likely to contain. The absence of a plateau in any curve indicates that additional deposits continue to introduce new named entities (e.g., new terms, locations, genes, and methods), see Figure 4. Across all repositories, this represents a substantial opportunity for growth; however, the growing number of different entities will likely include many near-synonyms, which could be reduced by curation to support the use of controlled vocabularies and ontologies. This would improve interoperability, even if the number of unique entities continues to increase Guizzardi (2020); Wilkinson et al. (2016).

Ontologies facilitate connections between multiple databases by providing a standardised vocabulary for a specific domain as implemented in IDR. When these terms are consistently included as part of the metadata vocabulary, they can be applied in user interfaces for databases that utilise ontology-based annotations and natural language processing methods Hoehndorf, Schofield, and Gkoutos (2015). Furthermore, ontologies can serve as links between different classes, enabling access to ontology-based annotations, such as those provided by GOPubMed Doms and Schroeder (2005), which allows users to retrieve scientific articles based on ontology classifications Hoehndorf et al. (2015).

Although structured metadata fields are essential for data organisation and retrieval, their number and specific content vary greatly, even among the three repositories examined in this study. Our analysis focuses primarily on free-text fields, omitting information contained in structured metadata. Free-text fields differ substantially across repositories in terms of schema, required elements, and use of controlled vocabularies. It is also worth noting that curated data is not necessarily reflected in the free-text fields. For example, information such as gene names or compound names may be recorded exclusively as key-value pair annotations, and may therefore be absent from the description fields altogether. Nevertheless, the flexibility of free-text fields enables meaningful comparisons across datasets.

The variability in requirements presents a significant challenge for biologists, who frequently interact with multiple repositories, each with its own set of requirements. Completing structured metadata fields is often a repetitive and time-consuming task, even considered mundane work Ribeiro, Meckin, Balmer, and Shapira (2023); Sambasivan, Kapania, and Highfll (2021), highlighting an important opportunity for natural language processing (NLP) and large language models (LLMs) to facilitate the extraction and standardisation of structured metadata from free-text descriptions.

Considering these findings, analyses such as those presented in this study show that richness metrics are not merely descriptive; they can also inform curation practices and inspire the design of submission forms. Furthermore, various stakeholders—including researchers, data users, and data managers—can rely on this framework to evaluate the status of their metadata.

By applying these straightforward metrics, different types of users can gain a clearer understanding of the information’s content and identify potential gaps or deficiencies in the metadata they use or produce. For instance, methodologies that quantify richness can be used for quality control to detect anomalous entity profiles and unexpectedly low information density within datasets. While entity recognition offers a valuable means of capturing the overall characteristics of a dataset, it can occasionally introduce discrepancies between the assigned category labels and the terminology extracted from unstructured text, thereby potentially compromising the accuracy of the results.

Future research should focus on the application of these methods, in combination with Large Language Models (LLMs), to improve existing metadata. These models can enhance quality by suggesting missing information, completing incomplete entries, and recommending additional categories, including standard metadata fields. This strategy should be explored across diverse types of repositories, particularly those where metadata is provided in free-text descriptions.

## 5 Acknowledgment

The authors gratefully acknowledge **Dr Jean-Marie Burel** (Molecular Cell and Developmental Biology, School of Biological Sciences, University of Dundee) for sharing valuable knowledge about the IDR metadata scheme and for his valuable observations on the article content, and **Dr Daniel Thèdie** for his diligence in providing BioDare2 metadata (BioRDM team, School of Biological Science, University of Edinburgh).

## 6 Authors contributions

**Maria Juliana Rodriguez-Cubillos:** Conceptualisation, Data Curation, Formal analysis, Investigation, Methodology, Software, Writing—original draft, Writing—review & editing. **Tomasz Zieliński:** Conceptualisation, Methodology, Supervision, Writing—review & editing. **Jason Swedlow:** Conceptualisation, Methodology, Supervision, Visualisation, Writing—review & editing. **T. Ian Simpson:** Conceptualisation, Methodology, Investigation, Formal analysis, Software, Supervision, Visualisation, Writing—review & editing. **Andrew J. Millar:** Conceptualisation, Formal analysis, Funding acquisition, Investigation, Methodology, Supervision, Project administration, Visualisation, Writing—review & editing.

## 9 Data availability

- **Datasets:** All the data used for the analysis of IDR and DataShare is publicly available in the correspondence database. The dataset used is publicly available in Zenodo (DOI 10.5281/zenodo.22229073): https://zenodo.org/records/22229074.

- *BioDare2* : https://biodare2.ed.ac.uk/
- *DataShare:* https://datashare.ed.ac.uk/rest
- *IDR*: https://github.com/IDR/idrmetadata.
- **Code:** The original code used for the analysis is available in the GitHub public repository: https://github.com/mjrodriguezc/metadata_project/tree/main/src/Repositories_analysis

## 10 Generative AI statement

The author(s) declared that generative AI was used in the creation of this manuscript. Generative AI was used for language editing and grammatical refinement of the manuscript. All AI-assisted content was reviewed, edited, and verified by the human authors.

## Supplementary Material

**Figure S1:**
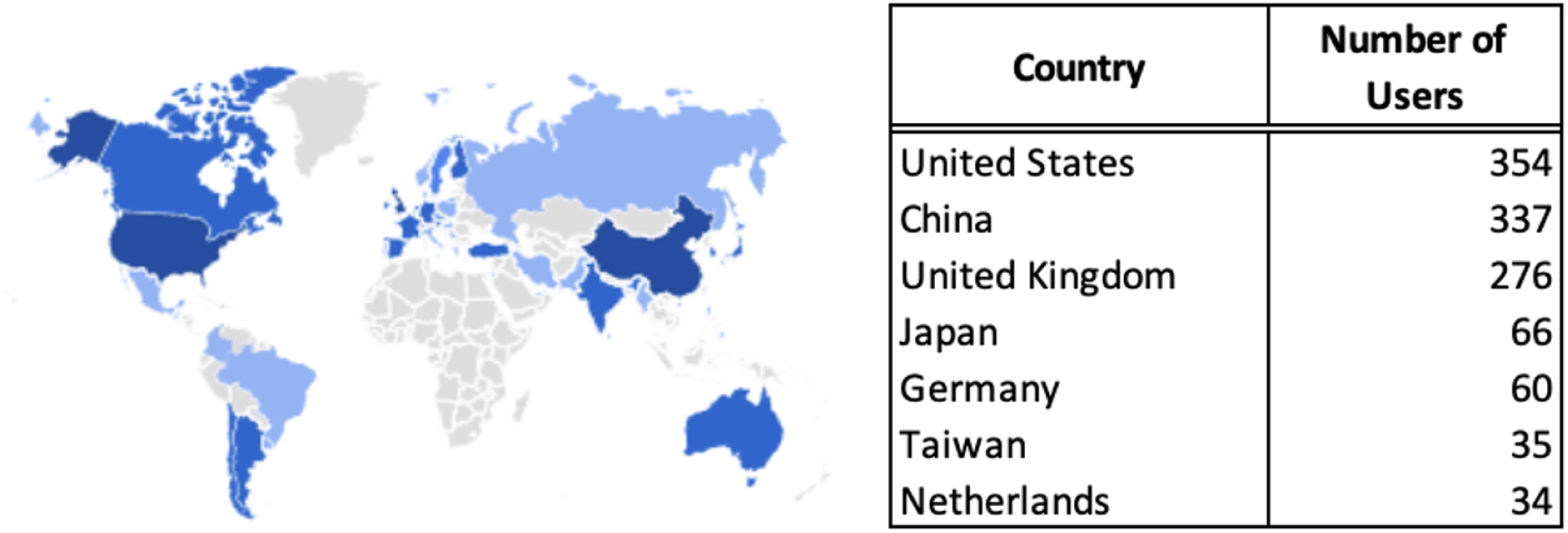
World representation by colour intensity of the number of users per country of the BioDare2 **?** from 01st January to November 17th of 2023. A darker blue represents more user density in the defined period **?**

**Figure S2:**
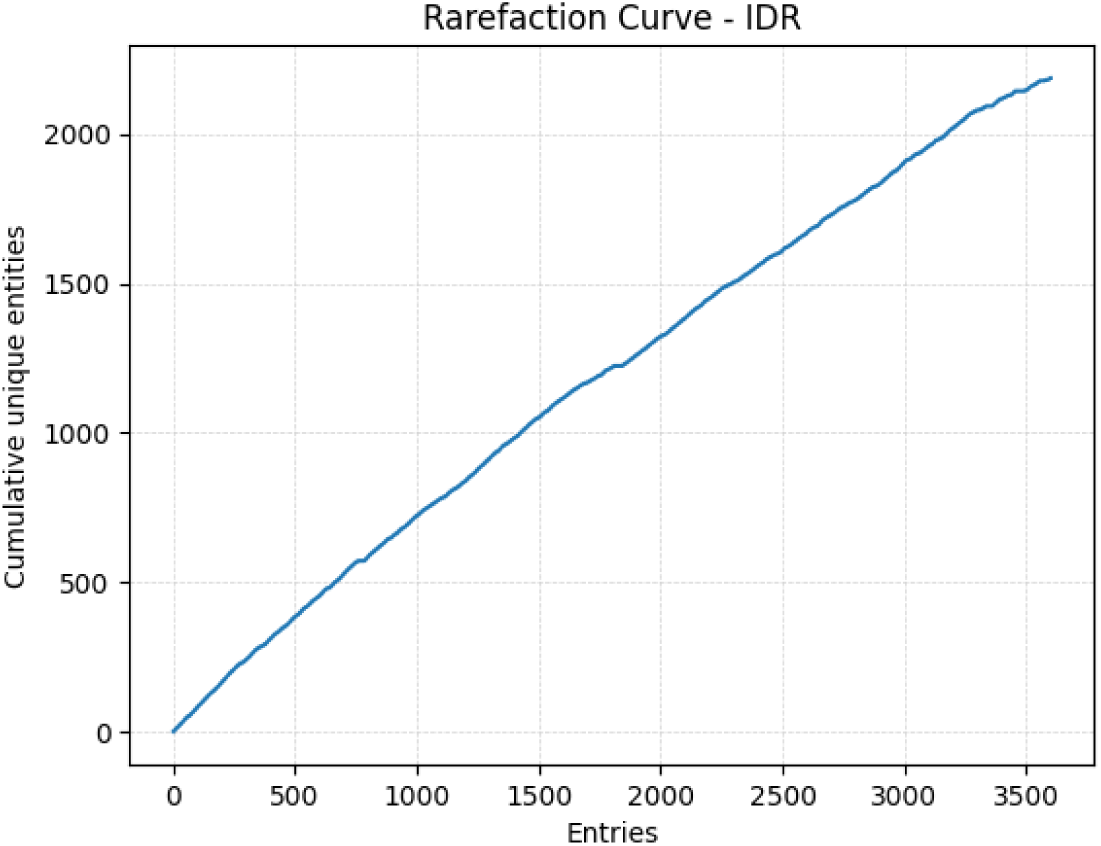
Rarefaction analysis of entity discovery with increasing entries. Rarefaction curves show the relationship between the number of entries sampled and the cumulative number of unique entities identified in IDR..

**Figure S3:**
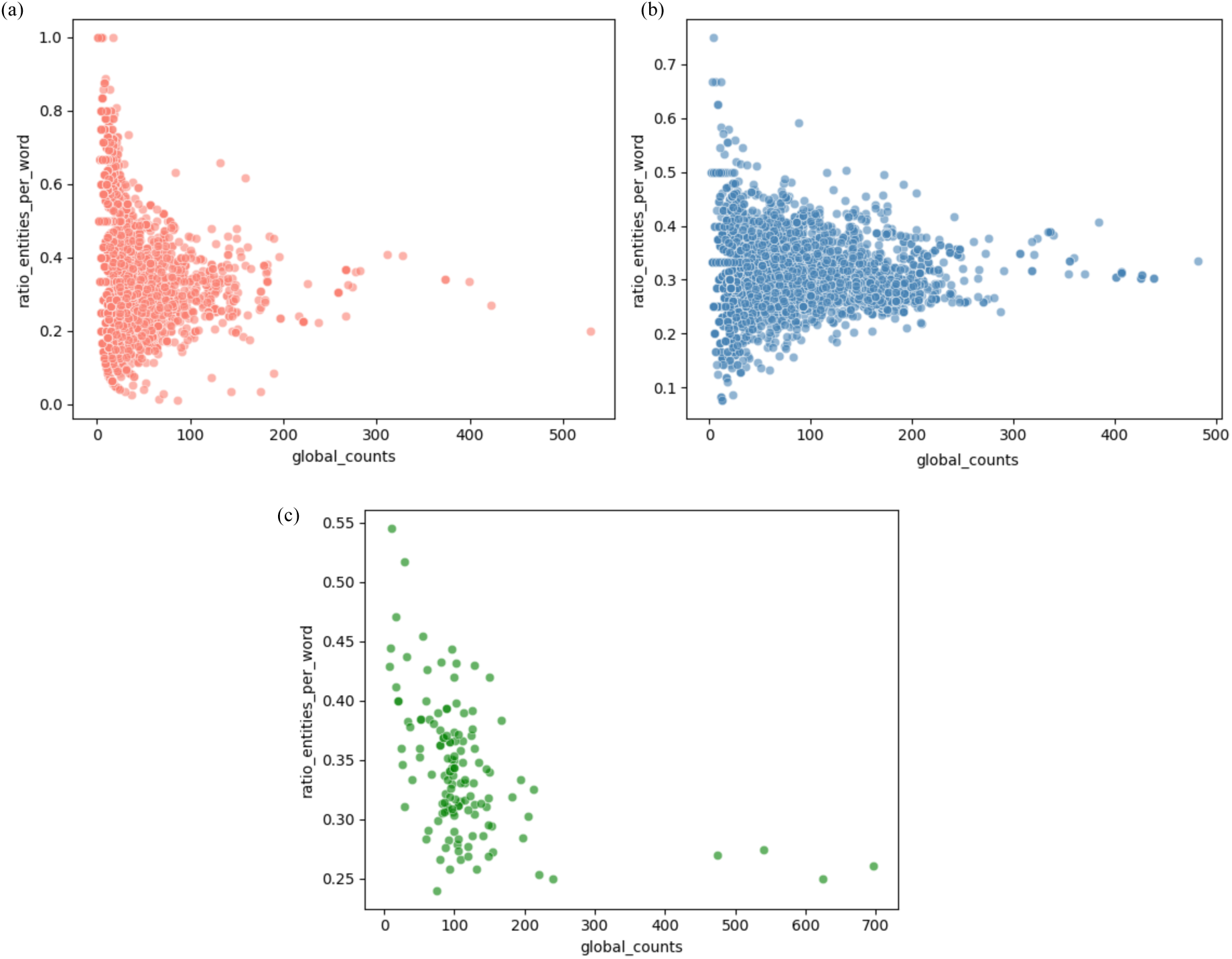
Scatter plot of the ratio of entities per word versus global counts for (a) BioDare2, (b) DataShare, and (c) IDR. Notably, for the first two repositories, the majority of entries are clustered between 0.2 and 0.5, whereas IDR exhibits a distinct distribution, with certain entries possessing a large number of words but a relatively small ratio of entities.

