## Supplementary Figure 1 for "The Shape of Biological Metadata: Measuring Repository Richness with Entity-Based NLP Metrics"

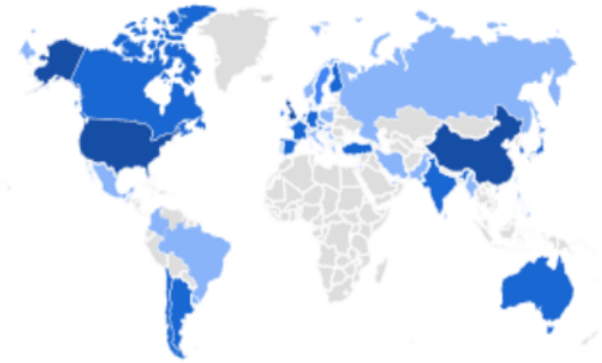

| Country | Number of Users |
|---|---|
| United States | 354 |
| China | 337 |
| United Kingdom | 276 |
| Japan | 66 |
| Germany | 60 |
| Taiwan | 35 |
| Netherlands | 34 |

**Supplementary Figure 1.** World representation by colour intensity of the number of users per country of the BioDare2 (Zielinski et al., 2022) from 01st January to November 17th of 2023. A darker blue represents more user density in the defined period (McMullen, 2010).
