## Supplementary Figure 2 for "The Shape of Biological Metadata: Measuring Repository Richness with Entity-Based NLP Metrics"

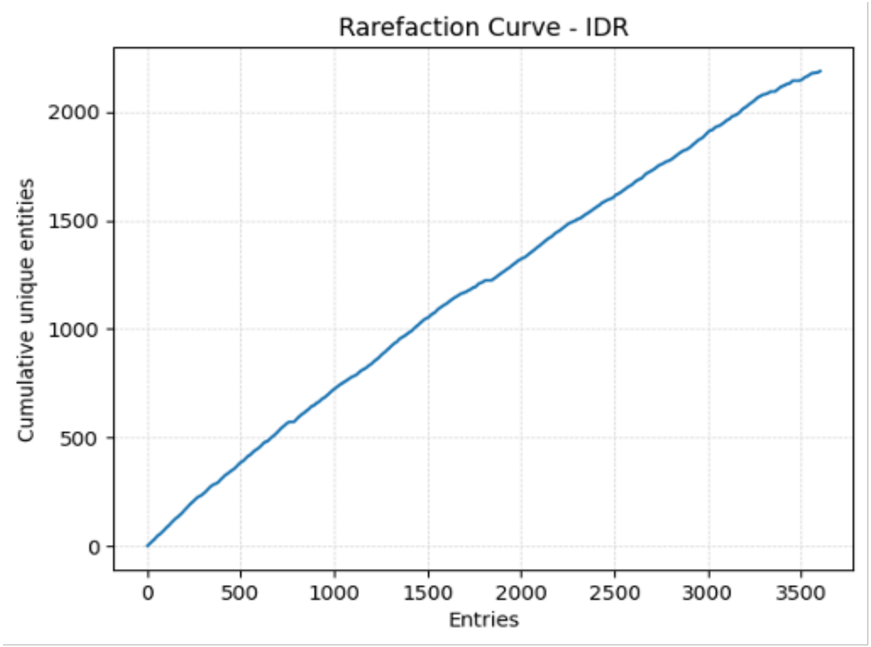


**Supplementary Figure 2.** Rarefaction analysis of entity discovery with increasing entries. Rarefaction curves show the relationship between the number of entries sampled and the cumulative number of unique entities identified in IDR.
