## Supplementary Figure 3 for "The Shape of Biological Metadata: Measuring Repository Richness with Entity-Based NLP Metrics"

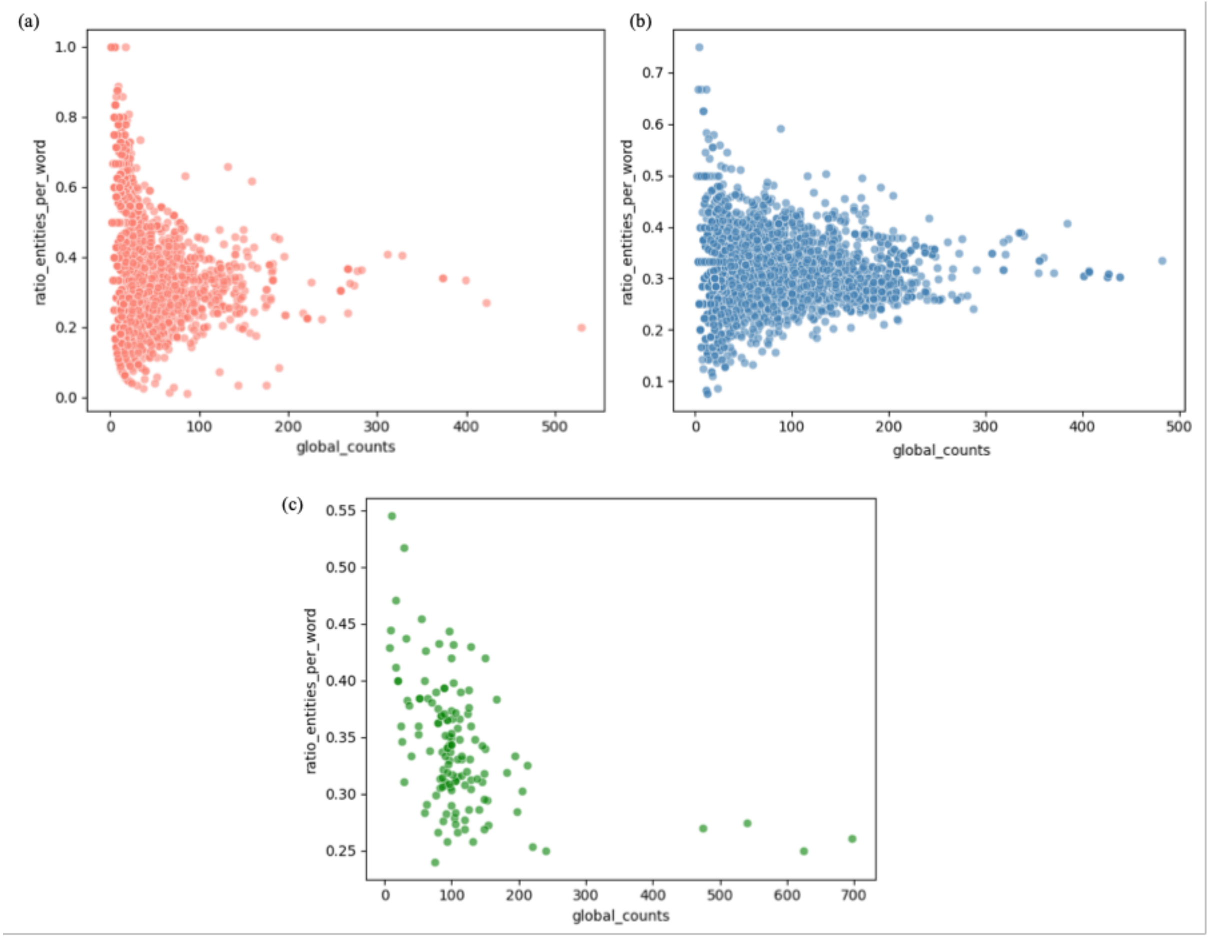


**Supplementary Figure 3.** Scatter plot of the ratio of entities per word versus global counts for (a) BioDare2, (b) DataShare, and (c) IDR. Notably, for the first two repositories, the majority of entries are clustered between 0.2 and 0.5, whereas IDR exhibits a distinct distribution, with certain entries possessing a large number of words but a relatively small ratio of entities.
